# Complex multicellularity in fungi: evolutionary convergence, single origin, or both?

**DOI:** 10.1101/230532

**Authors:** László G. Nagy, Krisztina Krizsán

**Author notes:** Correspondence should be addressed to L.G.N.

## Abstract

Complex multicellularity comprises the most advanced level of organization evolved on Earth. It has evolved only a few times in metazoans, green plants, brown and red algae and fungi. Compared to other lineages, the evolution of multicellularity in fungi follows different principles; both simple and complex multicellularity evolved via unique mechanisms not seen in other lineages. In this article we review ecological, paleontological, developmental and genomic aspects of complex multicellularity in fungi and discuss the general principles of the evolution of complex multicellularity in light of its fungal manifestations. Fungi represent the only lineage in which complex multicellularity shows signatures of convergent evolution: it appears 8-12 distinct fungal lineages, which show a patchy phylogenetic distribution, yet share some of the genetic mechanisms underlying complex multicellular development. To mechanistically explain the patchy distribution of complex multicellularity across the fungal tree of life we identify four key observations that need to be considered: the large number of apparently independent complex multicellular clades; the lack of documented phenotypic homology between these; the universal conservation of gene circuits regulating the onset of complex multicellular development; and the existence of clades in which the evolution of complex multicellularity is coupled with limited gene family diversification. We discuss how these patterns and known genetic aspects of fungal development can be reconciled with the genetic theory of convergent evolution to explain its pervasive occurrence in across the fungal tree of life.

## 1. Introduction: Simple and complex multicellularity

Multicellularity comes in many forms and complexity levels, ranging from simple cell aggregations, colonies, films or filaments to the most complex organisms known(Aguilar, Eichwald & Eberl, 2015; Fairclough, Dayel & King, 2010; Herron & Nedelcu, 2015; Knoll, 2011; Niklas, 2014; Niklas & Newman, 2013b; Rainey & de Monte, 2014; Richter & King, 2013; Rokas, 2008; Sebé-Pedrós, Irimia, del Campo *et al.*, 2013; Szathmary & Smith, 1995; Umen, 2014). While simple cell aggregations and colonies evolved at least 25 times in both pro- and eukaryotes(Grosberg & Strathmann, 2007; Rokas, 2008), complex multicellularity has evolved in up to five major groups, animals, embryophytes, red and brown algae(Claessen, Rozen, Kuipers *et al.*, 2014; Cock, Godfroy, Strittmatter *et al.*, 2015; Cock, Sterck, Rouze *et al.*, 2010; Knoll, 2011; Nagy; Niklas, 2014; Niklas & Newman, 2016; Sebe-Pedros, Degnan & Ruiz-Trillo, 2017; Umen, 2014), and fungi. While for many groups evolving simple multicellularity seems to be relatively easy, complex multicellularity probably represents a more difficult leap for organisms(Grosberg *et al.*, 2007). Simple and complex multicellularity are distinguished based on the proportion of cells being in direct contact with the environment (some vs. all), the extent of cellular differentiation, cell adhesion, communication, a developmental program and programmed cell death (PCD)(Cock *et al.*, 2010; Herron *et al.*, 2015; Knoll, 2011; Knoll & Hewitt, 2011). Complex multicellularity is usually used in reference to 3-dimensional differentiated structures, although how (and whether) it is defined varies widely across studies. Here, we focus on a genetically determined developmental program, determinate growth and 3dimensional organization as key traits for complex multicellularity. The rationale for this is that these traits represent major hurdles to evolving higher-level complex organization, that in 3dimensional structures not all cells are in direct contact with the environment, necessitating mechanisms for overcoming the limitations of diffusion and for cell-cell adhesion. On the other hand, primitive mechanisms for cell adhesion, communication and differentiation exist also in simple colonial and even unicellular protists(King, Hittinger & Carroll, 2003), even though they reach their highest complexity in complex multicellular organisms. Similarly, PCD occurs also in uni- and simple multicellular lineages(Herron *et al.*, 2015), so the more relevant question in the context of complex multicellularity is whether unprogrammed cell death is lethal to the multicellular individual or stalls its further development and reproduction(Knoll, 2011). It should be noted that, as is often the case in biology, discretely categorizing a continuum of evolved forms can be challenging, nevertheless, the distinction of simple and complex MC is useful for comparing phyletic and genetic patterns across distantly related multicellular groups.

The main focus of this review is the convergent evolution of complex multicellularity from a fungal perspective. We discuss how genetic and developmental information can be reconciled with the multiple origins of complex MC in fungi to understand its evolutionary history. We first demonstrate that complex MC is so widespread in fungi that it challenges our general view of its origins by convergent evolution. We then evaluate alternative hypotheses on the genetic mechanisms of the evolution of complex MC in fungi and how emerging theories of convergent evolution can inform our understanding of the evolution of MC. We start by introducing a concept for distinguishing simple and complex multicellular grades of fungal evolution, then discuss phylogenetic, developmental and genetic aspects of complex multicellularity in the fungal world.

## 2. Simple multicellularity in fungi

Multicellular organisms have diverse unicellular ancestry. Presumably, most multicellular eukaryotes evolved from aggregative or colony-forming ancestors, resembling extant choanoflagellates(Fairclough *et al.*, 2010; Hanschen, Marriage, Ferris *et al.*, 2016; Richter *et al.*, 2013; Sebe-Pedros *et al.*, 2017) and volvocine algae, among others(Fairclough *et al.*, 2010; Hanschen *et al.*, 2016; King, 2004; Niklas, 2014; Niklas *et al.*, 2016; Richter *et al.*, 2013; Rokas, 2008; Sebe-Pedros *et al.*, 2017; Telford, Budd & Philippe). Here, the evolution of sophisticated mechanisms for cell adhesion and cell-cell communication followed by functional and morphological differentiation defines the 'classic' route to multicellularity(Brunet & King, 2017). The evolution of multicellularity in fungi departs from this classic scheme in many respects. Fungi develop multicellular thalli composed of hyphae that extend apically and grow and branch under rules similar to fractal geometry. Hyphae most likely evolved for optimizing foraging efficiency; they direct growth and occupy space to maximize substrate utilization, resulting in a loosely arranged, interconnected, fractal-like network. Hyphae are hypothesized to have evolved by the gradual elongation of substrate-anchoring rhizoids of unicellular ancestors resembling extant Chytridiomycota(Harris, 2011), although alternative routes in convergently evolved hyphal forms (e.g. Monoblepharidomycetes(Dee, Mollicone, Longcore *et al.*, 2015)) may exist. Nevertheless, the evolution of fungal hyphae likely did not involve the modification of cell wall biogenesis for daughter cells to remain together, as seen in filamentous bacteria and algae(Claessen *et al.*, 2014; Herrero, Stavans & Flores, 2016; Niklas, 2014) or snowflake yeast(Ratcliff, Denison, Borrello *et al.*, 2012; Ratcliff, Herron, Howell *et al.*, 2013). The first hyphae were probably similar to those of extant Mucoromycota and gradually evolved sophisticated mechanisms for septum formation, nutrient and organelle trafficking, branch site selection, etc (for recent reviews on hyphal morphogenesis see (Harris, 2011; Lew, 2011)). Primitive hyphae were uncompartmentarized coenocytic multinucleate structures where the free flow of cell content was probably hardly regulated. In modern hyphae, hyphal segments are closed off from the growing tip by septa and various septal occlusures, such as Woronin bodies, dolipores or simpler amorphous materials.

Thus, simple MC in fungi likely evolved via a linear process that could have avoided some of the hurdles that should be overcome for establishing an evolutionarily stable multicellular organization(Brown, Kolisko, Silberman *et al.*; Du, Kawabe, Schilde *et al.*, 2015). Hyphae might not face group conflicts and could bypass the need for fitness alignment between individual cells to directly confer a higher exported organism-level fitness, or handle conflicts at the level of individual nuclei. Fractal-like filling of the available space might further minimize conflict among separate hyphae of the same individual. Similar ‘siphonous->multicellular’ transformations can be found in certain algae(Niklas, Cobb & Crawford, 2013a; Niklas *et al.*, 2013b) and may represent a third way to evolve simple multicellularity in addition to the colonial and aggregative routes(Brown *et al.*; Brunet *et al.*, 2017; Sebé-Pedrós *et al.*, 2013).

However, fungal mycelia do not show all characteristics of complex MC. The growth of vegetative mycelia is indeterminate and cellular differentiation is mostly limited to asexual or sexual spores (conidia, zygo-, asco- and basidiospores, etc.) and cells involved in (a)sexual reproduction and not spatially or temporally integrated into a developmental program. Further, all cells are in direct contact with the external environment, which means that nutrient and O_2_ uptake through diffusion is not impeded by a compact, 3-dimensional organization. Although programmed cell death is widely observed, unprogrammed cell death is not lethal to the entire organism. Thus, we consider vegetative mycelia as a grade of simple multicellularity, with noting that some species' vegetative mycelia are capable of complex functionalities and can differentiate several distinct cell types.

## 3. Complex multicellularity in fungi

We here define complex multicellularity as structures showing a 3-dimensional differentiated organization with a spatially and temporally integrated developmental program that grows until reaching a genetically predetermined shape and size. Complex MC in fungi is mostly discussed in the context of sexual fruiting bodies (Fig 1), although fungi produce a plethora of other complex multicellular structures, such as asexual fruiting bodies, rhizomorphs, mycorrhizae or sclerotia (Fig 2, see below). Fruiting bodies are 3-dimensional structures that enclose reproductive cells and the developing spores into a protective environment and facilitate spore dispersal both passively and actively(Dressaire, Yamada, Song *et al.*, 2016; Roper, Seminara, Bandi *et al.*, 2010). This immediately highlights the most crucial difference between fungi and other complex MC organisms. Whereas in other lineages complex MC comprises the reproducing individual, it refers to specific structure(s) of the fungal individual. Complex multicellularity in fungi fulfills mostly reproductive roles, whereas for feeding through osmotrophy, foraging for nutrients and exploration of the substrate simple multicellularity clearly represents a better adaptation. Simple and complex MC coexist in the same species in fungi. Fruiting body forming fungi undergo a transition from simple to complex multicellularity as part of their life cycle, which not only makes them unique among complex multicellular organisms, but also a potentially useful model system to study complex multicellular development.

**Figure 1.**
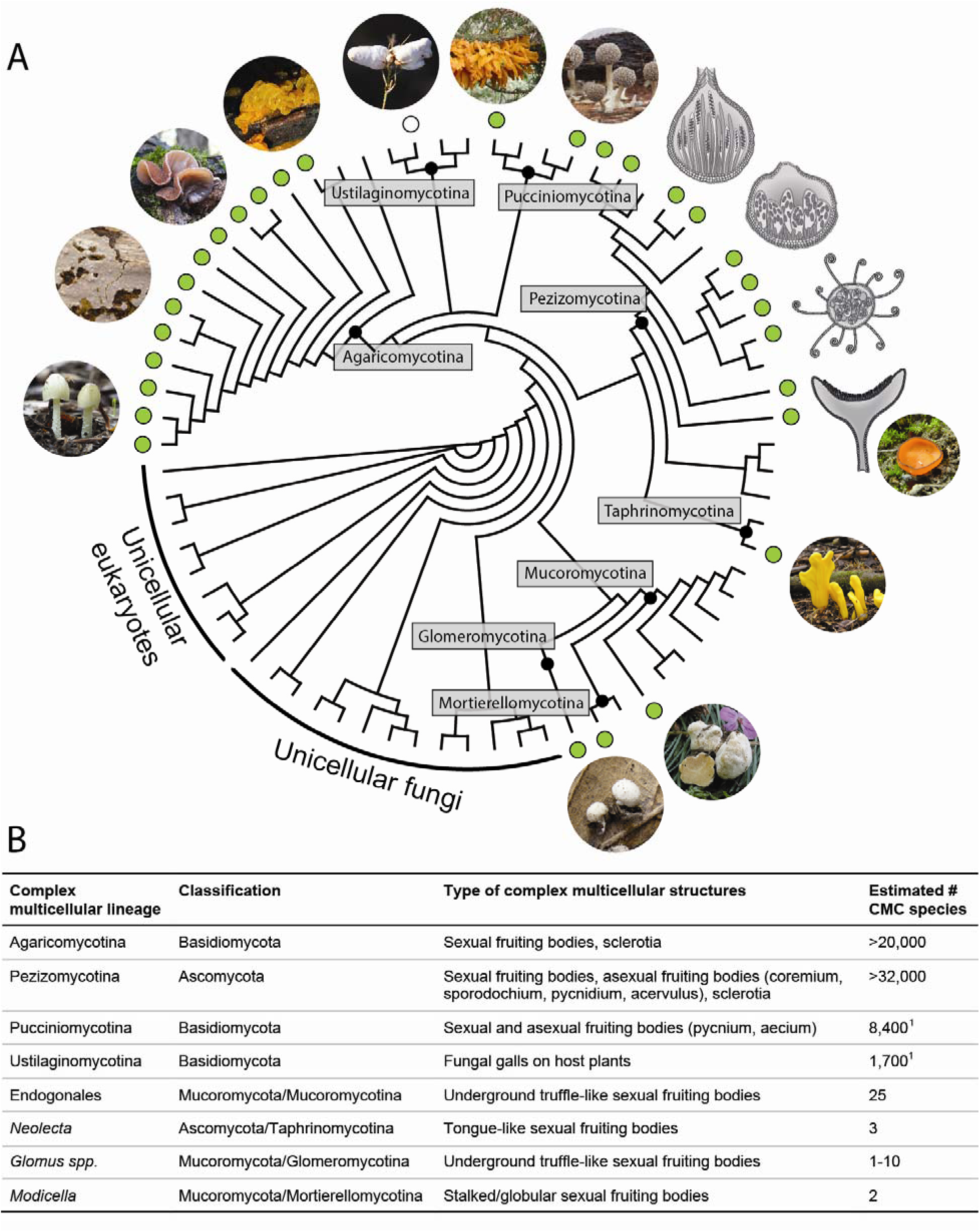
The phylogenetic distribution of complex multicellularity in fungi. **A.** Most typical complex multicellular morphologies of sexual fruiting bodies are shown for each major clade of fungi. Species pictured from left to right are: *Bolbitius titubans, Gloeocystidiellum sp, Auricularia auricula-judae, Tremella mesenterica, Testicularia cyperi, Gymnosporangium clavariiforme, Phleogena faginea, Podospora anserina* (perithecium), *Mycosphaerella* sp. (pseudothecium), *Microspheara* sp. (cleisthothecium), *Peziza* sp (apothecium), *Neolecta irregularis, Endogone flammirocona, Modicella reniformis*. Green dots indicate lineages with known complex multicellular representatives; an empty circle at the Ustilaginomycotina refers to the uncertain status of the galls produced by *Testicularia* and allies. **B.** Classification, types of complex multicellular structures produced and estimated number of complex multicellular species for each major lineage of complex multicellular fungi. See acknowledgements for sources of images.

**Figure 2.**
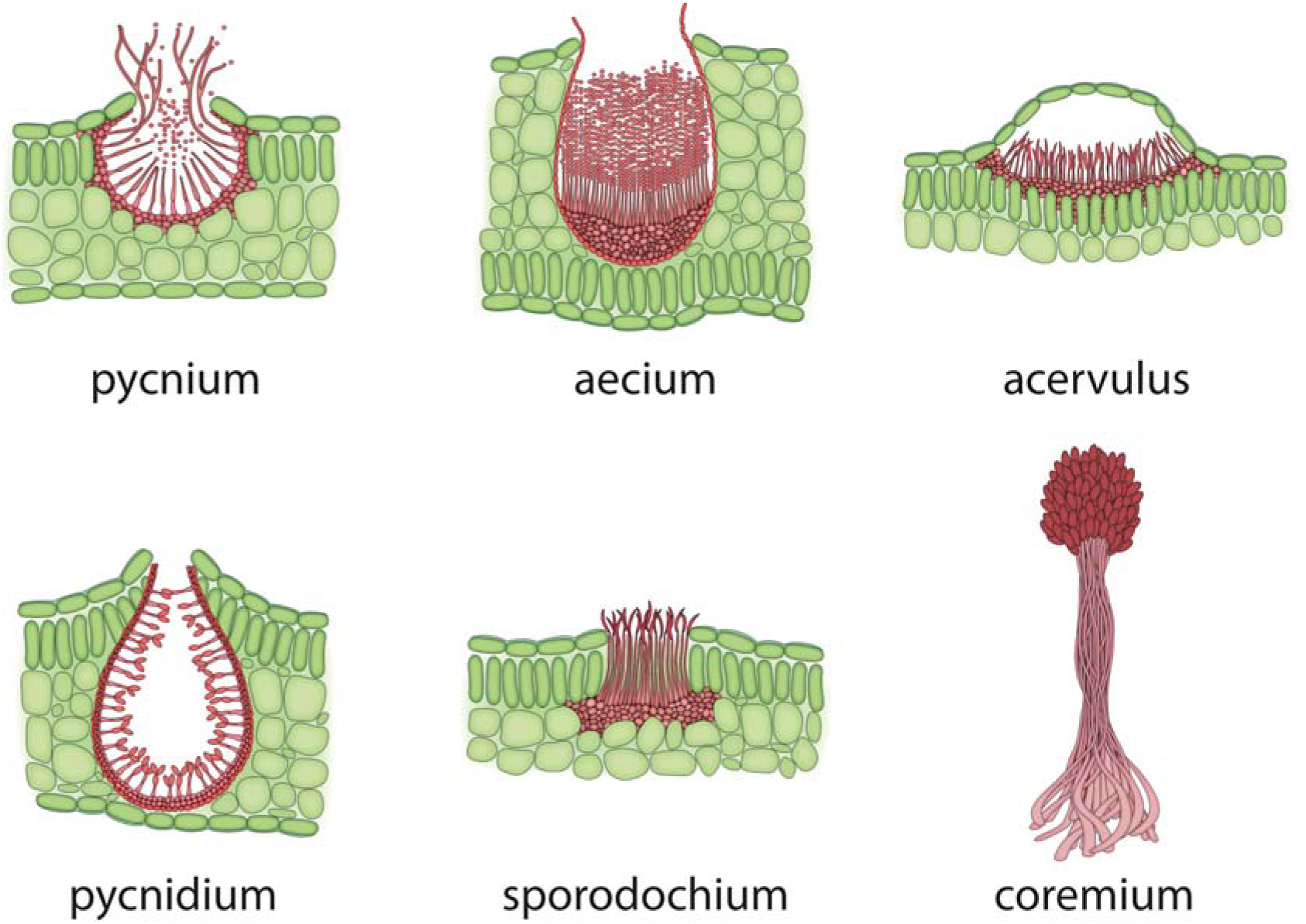
Asexual complex multicellular structures produced by fungi in the Pucciniomycotina (pycnium, aecium) and Ascomycota (acervulus, pycnidium, sporodochium, coremium = synnemata). Note that these structures, like all complex multicellular structures, have a genetically determined shape and size and a tighly integrated developmental program. Plant tissue shaded green.

Another important difference is that growth remains polarized in fruiting bodies, i.e. complex multicellular structures and organs therein are formed by the aggregation, elongation and specialization of hyphae, which has implication for the evolution of complex MC. For example, there is no need for a qualitatively new mechanism for long distance distribution of nutrients or O_2_(Woolston, Schlagnhaufer, Wilkinson *et al.*, 2011), as seen in complex animals and plants. It should be noted, though, it has been hypothesized that in the most complex fruiting bodies of Basidiomycota air channels are being formed by the deposition of hydrophobins along the cell walls.

What is the driving force for evolving complex MC in fungi? Avoiding predation has been named as one of the factors driving the evolution of increasingly complex and larger animals(Kaiser, 2001; Knoll, 2011; Rokas, 2008). This is, however, quite unlikely as a driving force in fungi, because of their osmotrophic lifestyle and because hyphal multicellularity is sufficient to avoid the entire individual being eaten by predators. Even if much of the thallus is destroyed, the individual can completely regenerate, as long as sufficient nutrients are available(Fricker, Heaton, Jones *et al.*, 2017). The most evident selective advantage for fruiting bodies is the promotion of spore dispersal and enclosure of developing sexual propagules into a 3-dimensional structure. Remarkably, both major sporogenous cell types, asci and basidia, evolved mechanisms for active spore discharge(Dressaire *et al.*, 2016; Roper *et al.*, 2010; Trail, 2007). Increasing the efficiency of spore dispersal could have driven the evolution of structures that enclose and raise asci and basidia above ground level. Fruiting bodies also provide protection against infections and predation of spores, through various structural (veils, setae, hairs, spikes) and chemical defense systems. Insecticidal and antimicrobial armories are particularly rich in fruiting-body forming fungi and include secondary metabolites, pore forming toxins(Plaza, Lin, van der Velden *et al.*, 2014), lectins, many of which are encoded by genes acquired horizontally from bacteria(Kunzler, 2015).

In addition to sexual fruiting bodies, fungi produce a plethora of structures that conform to some to all aspects of complex multicellularity. Asexual fruiting bodies of Ascomycota (pycnidia, acervuli, sporodochia and coremia) are three dimensional reproductive structures, that harbor asexual spores (conidia). They range from submacroscopic sizes to several centimeters (Figure 2) and are made of more or less tightly arranged hyphae. Size, shape and coloration are genetically determined, but cellular differentiation is often limited to a few cell types. Pycnia, uredinia, aecia, telia (including macroscopic telial horns) are reproductive structures of rust fungi (Pucciniomycotina). Although mostly sub-macroscopic (except telial horns of *Gymnosporangium spp*), they have a predetermined developmental program and show cell differentiation and adhesion of almost isodiametric cells (Figure 2). Ectomycorrhizae, rhizomorphs and sclerotia are also three-dimensional structures(Kues, 2000), however, there might be a looser genetic control over their size and shape.

## 4. Convergent origins of complex multicellularity in fungi

Complex multicellularity evolved in only five eukaryotic groups(Brawley, Blouin, Ficko-Blean *et al.*, 2017; Cock *et al.*, 2010; Niklas, 2014; Parfrey, Lahr, Knoll *et al.*, 2011; Umen, 2014). Within fungi, it occurs in most major clades and shows signs of convergent evolution(Knoll, 2011; Schoch, Sung, López-Giráldez *et al.*, 2009; Sugiyama, Hosaka & Suh, 2006; Taylor & Ellison, 2010) (Fig 1). The best known complex multicellular clades are the Pezizomycotina and the Agaricomycotina in the Asco- and Basidiomycota, respectively, where the majority fruiting body forming fungi belong (Fig 1). Although generally two origins of complex multicellularity are mentioned in fungi(Knoll & Lahr, 2016; Schoch *et al.*, 2009; Stajich, Berbee, Blackwell *et al.*, 2009), complex multicellular structures also occur in the early diverging Mucoromycota, the primarily yeast-like Taphrinomycotina as well as the Puccinio- and Ustilaginomycotina. Of these, the earliest diverging is the Mucoromycota, which primarily contains simple multicellular molds but also three small groups of fruiting body forming fungi. Members of the Endogonales (Mucoromycotina) form globose, truffle-like sporocarps filled with zygospores (Fig 1), while *Modicella* (Mortierellomycotina) forms small stalked fruiting bodies that contain sporangia and sporangiospores(Smith, Gryganskyi, Bonito *et al.*, 2013) (Fig 1). Similarly, several Glomeromycotina species produce small, underground sporocarps(Smith *et al.*, 2013). The Taphrinomycotina (Ascomycota) contains a single known fruiting body forming genus, *Neolecta* that forms brightly colored irregular or tongue-like fruiting bodies on soil(Nguyen, Cisse, Yun Wong *et al.*, 2017). This genus is particularly interesting from a developmental perspective as it is nested in a clade of primarily unicellular yeasts, and has a yeast-like genome architecture(Nagy; Nguyen *et al.*, 2017).

In the Basidiomycota, the largest fruiting body producing lineage is the Agaricomycotina, where multicellularity reached its highest complexity in fungi. Nearly all of the species produce fruiting bodies, with the exception of some secondarily reduced yeast lineages in the Tremellomycetes(Nagy, Ohm, Kovács *et al.*, 2014) or ant-associated Pterulaceae that might have lost the ability to form fruiting bodies(Mueller, 2002). This group also includes the most typical manifestations of agaricoid 'mushroom' morphologies as well as an array of morphologically diverse forms(Hibbett, 2007; Hibbett & Binder, 2002). Aside from the Agaricomycotina, complex multicellular species are found in the Puccinio- and Ustilaginomycotina (rust and smut fungi, respectively) as well, although they comprise the minority of species in their clades compared to simple or yeast-like species. Fruiting bodies are known in at least 4 classes of the Pucciniomycotina(Aime, Matheny, Henk *et al.*, 2006) (Atractelliomycetes, Agaricostilbomycetes, Pucciniomycetes, Microbotryomycetes) and include simple capitate (*Phleogena*) or cup-shaped (*Platygloea, Kriegeria, Fig 1)* morphologies, but also crust-like (e.g. *Septobasidium*) and gelatinous (e.g. *Helicogloea*) forms resembling those found in early-diverging Agaricomycotina. As the relationships of fruiting body forming classes of the Pucciniomycotina are still unresolved(Aime *et al.*, 2006; Bauer, Begerow, Sampaio *et al.*, 2006; Wang, Groenewald, Takashima *et al.*, 2015), there is some uncertainty as to the number of independent origins of fruiting body development in this subphylum. The occurrence of true fruiting bodies in the Ustilaginomycotina may be controversial. *Ustilago maydis* was recently reported to produce fruiting body-like structures in vitro(Cabrera-Ponce, Leon-Ramirez, Verver-Vargas *et al.*, 2012) whereas other species (e.g. *Testicularia* spp., *Exobasidium* spp.) produce gall-like swellings on parasitized plants that are mostly made up of fungal hyphae but incorporate more or less of the plant tissue too. Although these show some features of complex multicellularity (e.g. tight arrangement of hyphae, adhesion), whether their development follows a genetically pre-determined program or their growth is determinate, remain to be understood(Nagy).

The phylogenetic distribution of complex multicellular fungi is patchy and the above mentioned lineages outline at least 8 complex multicellular clades. However, the Pucciniomycotina, Glomeromycota and potentially the Ustilaginomycotina may comprise more than a single origin of fruiting body producing species, yielding 12 as a conservative upper estimate for the number of independent complex multicellular clades in fungi, although this may need refinement as more resolved phylogenies become available.

## 5. Evolutionary timescale for complex multicellular fungi

Complex multicellular organisms are of vastly different ages, yet their origins and diversification might have required some basic geologic formations, eukaryotic prehistory and abiotic conditions (e.g. O_2_ or sulfide concentrations(Canfield, Poulton & Narbonne, 2007; Canfield & Teske, 1996; Johnston, Poulton, Dehler *et al.*, 2010; Richter *et al.*, 2013)). Whereas simple multicellular lineages can be as old as 3.5 Ga(Aguilar *et al.*, 2015), complex multicellular organisms originated much later. Recent studies(dos Reis, Thawornwattana, Angelis *et al.*, 2015; Parfrey *et al.*, 2011; Sharpe, Eme, Brown *et al.*, 2015) dated complex MC clades between 175 to 800 myr, with the Metazoa being the oldest (700-800 myr), followed by Florideophyceae red algae(Parfrey *et al.*, 2011; Xiao, Knoll, Yuan *et al.*, 2004) (550-720 myr), the Embryophyta (430-450 myr) and macroscopic brown algae (175 myr)(Silberfeld, Leigh, Verbruggen *et al.*, 2010). Due to the soft texture of fungal fruiting bodies, the fossil record is very patchy and available fossilized fruiting bodies are way too recent to provide reasonable estimates for the age of CMC clades(Berbee & Taylor, 2010; Cai, Leschen, Hibbett *et al.*, 2017; Hibbett, Grimaldi & Donoghue, 1997; Hibbett, Binder, Wang *et al.*, 2003; Hibbett, Grimaldi & Donoghue, 1995; Poinar & Singer, 1990). Yet, the oldest known fruiting body fossil, a perithecium known as *Paleopyrenomycites devonicus*(*Taylor, Hass, Kerp et al., 2005*) is from the early Devonian (ca. 400 myr) placing the earliest physical evidence for complex multicellular Ascomycota roughly in the same age as the origin of embryophytes or red algae. Based on this, and other Ascomycota fossils, molecular clock analyses inferred the origins of the Pezizomycotina, the largest CMC clade in fungi, at 537 (443–695) myr(Beimforde, Feldberg, Nylinder *et al.*, 2014; Prieto & Wedin, 2013). The age of the Agaricomycotina have been inferred at 429-436 myr based on multiple calibration points and phylogenomic datasets(Chang, Wang, Sekimoto *et al.*, 2015; Floudas, Binder, Riley *et al.*, 2012; Kohler, Kuo, Nagy *et al.*, 2015). To our best knowledge, no molecular age estimates are available for *Endogone* and *Modicella*, nevertheless, their limited diversity and recent divergence from simple multicellular fungi suggest they are much younger than either the Agarico- or Pezizomycotina. Similarly, although chronological information is lacking for complex multicellular Puccinio-, Ustilaginomycotina and Taphrinomycotina, the patchy distribution of complex MC taxa in these clades suggests relatively recent origins. Taken together, the origins of the Pezizomycotina and Agaricomycotina seem to coincide with the origins of complex multicellular plants and algae in the Paleozoic, although significantly older estimates have, however, also been published(Berbee *et al.*, 2010; Heckman, Geiser, Eidell *et al.*, 2001). The much younger ages for smaller complex multicellular clades suggests that the evolution of complex MC in fungi is not tied to specific geologic events, as suggested for animals(Rokas, 2008), but was probably dependent on internal contingencies.

## 6. Complex multicellular functioning in fungi

How complex multicellularity manifests during fruiting body development has been of interest among mycologists for a long time. Information based on mutant screens and classical genetic techniques (Kues, 2000; Pöggeler, Nowrousian & Kück, 2006) is being increasingly complemented by high throughput studies based on whole genomes sequencing and RNA-Seq (Nowrousian, 2014). Studies involving whole transcriptome comparisons (Table 2) have revealed important principles of fruiting body development in both model and non-model fungal species. It is becoming evident, that in terms of morphogenesis and function, there are also a number of similarities and differences between fungi and other complex MC clades. In the following sections we therefore discuss known patterns of development, cell adhesion and signaling in complex multicellular fungi, with special emphasis on the general principles.

**Table 2.**
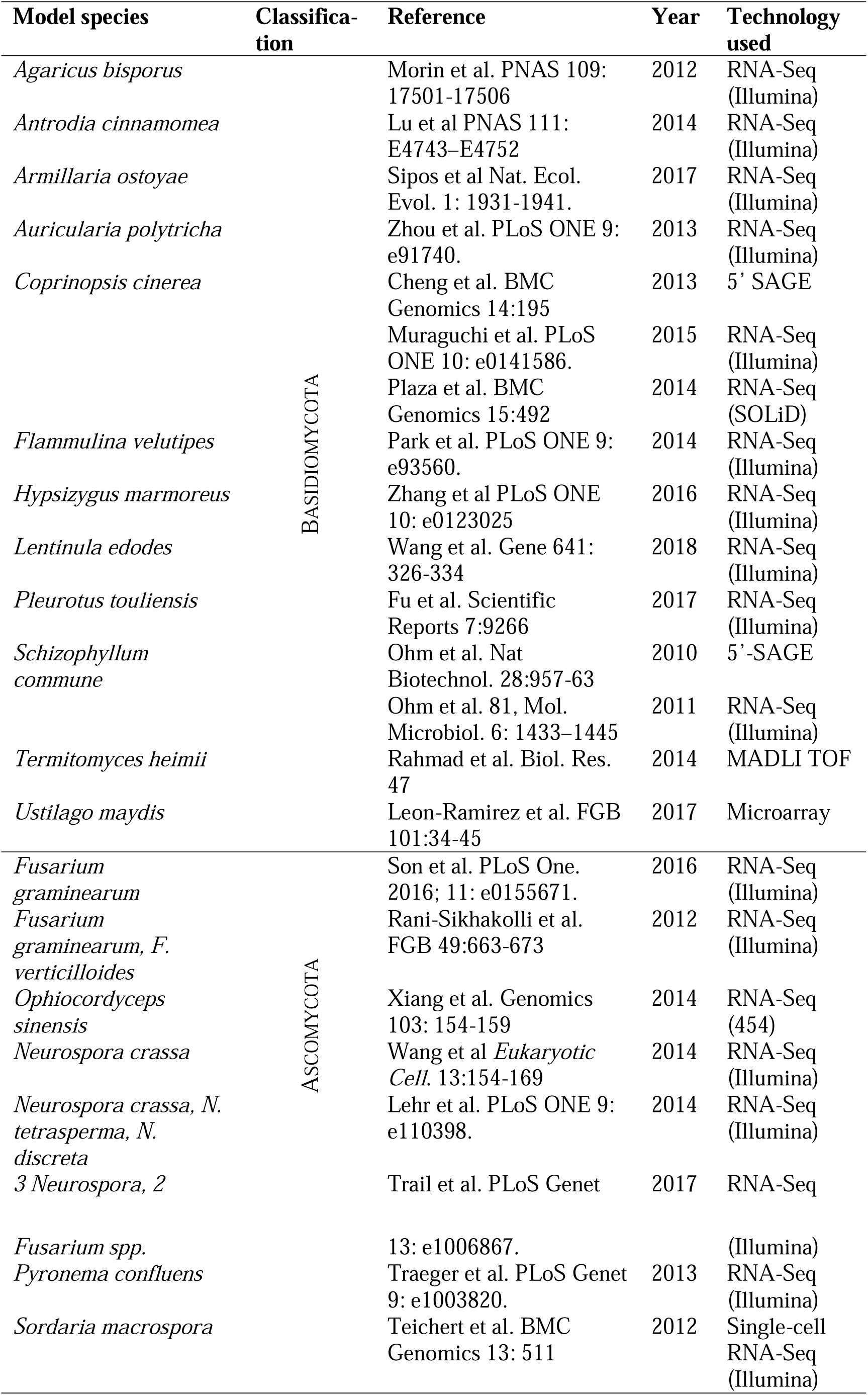
Published high-throughput gene expression studies of fungal multicellular development

### 6.1 Fungal development

Fungi are unique among complex multicellular organisms in that they can switch between simple and complex multicellularity during their life cycle. While the vegetative mycelium is composed of indeterminately growing hyphae that rarely adhere to each other, fruiting body development is a genetically predetermined process that involves adhesion, cell differentiation, growth, programmed cell death and senescence. It starts with a transition from a fractal-like growing vegetative mycelium to a 3-dimensional hyphal aggregate through intense localized hyphal branching and adhesion(Kues, 2000; Lakkireddy, Navarro-GonzÁLez, Velagapudi *et al.*, 2011; Lichius, Lord, Jeffree *et al.*, 2012; Pöggeler *et al.*, 2006) (Fig. 4). In the Basidiomycota, this aggregate is known as the primary hyphal knot. In the Ascomycota, development has been most widely studied in perithecium-forming Sordariomycetes (e.g. *Sordaria, Neurospora*), where the earliest complex multicellular stage is the protoperithecium (Fig. 4). The development of the hyphal knot and the protoperithecium involves the reprogramming of hyphal branching patterns to form the first step of complex MC. Subsequently, the differentiation of major tissue types takes place in secondary hyphal knots and perithecia in the Basidio- and Ascomycota, respectively. It has been estimated that perithecia can differentiate up to 13 cell tyes(Lord & Read, 2011), respectively, although the actual number of cell types, especially in the Basidiomycota, might be significantly higher.

The development of mature fruiting bodies follows genetically encoded programs that determine the species-specific morphologies(Kamada, 2002; Kamada, Sano, Nakazawa *et al.*, 2010; Kues, 2000; Pöggeler *et al.*, 2006; Trail & D.M., 2014), followed by senescence through the action of various oxidative (laccases, phenol oxidases), cell-wall degrading enzymes and tyrosinases, among others(Moore, 2005; Sakamoto, Nakade, Konno *et al.*, 2017). This is similar to death in other complex multicellular lineages and putatively serves the purpose of giving way to new reproducing generations of fruiting bodies and possibly recycling of cellular components towards reproductive cells(Moore, 2005). Growth remains apical even within fruiting bodies, but cell shape is extensively modified, ranging from hyphal to inflated and even isodiametric or polyhedral (referred to as conglutinate cells in *Sordaria(Lord et al., 2011)*), similar to animal and plant cells. Nonterminal cells might form side-branches, but regions of cell proliferation, resembling that in animals, to our best knowledge do not exist. Following a wave of cell differentiation events, growth in fruiting bodies is achieved by manipulating cell size through turgor and cell wall expansion.

There is evidence for autophagic cell death playing a role in sculpting fruiting bodies of both the Asco- and Basidiomycota. It should be noted that PCD of non-terminal cells may be counter-selected in fruiting body development, because it disrupts nutrient transport along the hyphae. Nevertheless, PCD has been reported to play a role in forming the gill cavity of *Agaricus bisporus(Lu, 2006; Umar & van Griensven, 1998)* (although this has been disputed) and in removing paraphyses from within ascomycete perithecia, presumably to give way to asci and spore release(Trail *et al.*, 2014). Further, autophagy genes are required for fruiting body development in *Sordaria macrospora(Voigt, Herzog, Jakobshagen et al., 2013; Voigt & Pöggeler, 2013)*, although how their defects disrupt development is not known yet.

### 6.2 Cell adhesion in fungi

Most of our knowledge on adhesive proteins of fungi pertains to adhesion to animal and plant hosts and various surfaces (e.g. medical devices) and comes primarily from simple multicellular and secondarily unicellular (i.e. yeast) species. Adhesion is mediated by a combination of sticky cell wall proteins and secreted carbohydrates, although the precise composition of fungal adhesives is very heterogeneous(de Groot, Bader, de Boer *et al.*, 2013; Epstein & Nicholson, 2016; Tucker & Talbot, 2001). Most cell wall proteins with glycosylphosphatidylinositol (GPI) anchoring(de Groot *et al.*, 2013; Sundstrom, 2002) to the cell wall have adhesive properties(Dranginis, Rauceo, Coronado *et al.*, 2007; Weig, Jansch, Gross *et al.*, 2004) and include adhesins(Sundstrom, 1999; Sundstrom, 2002; Weig *et al.*, 2004), flocculins(Dranginis *et al.*, 2007) and sexual agglutinins(Lipke & Kurjan, 1992) that participate in the adhesion of yeast cells to each other and to various surfaces. Other adhesive molecules include glycoproteins(Newey, Caten & Green, 2007) that are linked to cell wall sugars through N- or O-linked oligosaccharides(Bowman & Free, 2006) (mostly mannose or galactomannan) and secreted mannosyl and glucosyl residues. Although not much is known about the role and composition of the extracellular matrix in complex multicellular fungi, ECM deposition has been observed already in the earliest stages of fruiting body development(Lichius *et al.*, 2012).

Our understanding of cell adhesion within fruiting bodies is far from complete(Lord *et al.*, 2011; Trail *et al.*, 2014), nevertheless, many of the adhesive proteins described from simple multicellular fungi have been detected in fruiting bodies. GPI-anchored and fasciclin-like proteins(Liu, Chen, Min *et al.*, 2009; Miyazaki, Kaneko, Sunagawa *et al.*, 2007) have been implicated in cell adhesion within fruiting bodies(Trail, 2013), whereas hydrophobins have been suggested to form air channels and thus could help circumventing the limits of diffusion in 3-dimensional structures(Lugones, Wösten, Birkenkamp *et al.*, 1999). Similarly, lectins have been detected in fruiting bodies of Asco- and Basidiomycota, and might be involved in cell adhesion(Hassan, Rouf, Tiralongo *et al.*, 2015) but also in defense against predators(Hassan *et al.*, 2015). These families are conserved across the Asco- and Basidiomycota at the family level which consistent with both vertical inheritance of function and their parallel co-option for hypha-hypha adhesion in complex MC lineages. This would parallel adhesive molecules of complex animals having presumably evolved early in protist evolution for prey capture and later co-opted for cell-cell adhesion(Abedin & King, 2008; Abedin & King, 2010; King *et al.*, 2003; Richter *et al.*, 2013; Rokas, 2008).

### 6.3 Cell-cell communication and signaling

Multicellular organisms mediate transcriptional responses to external stimuli and synchronize cell functioning by various signal transduction pathways both within and between cells(de Mendoza, Sebé-Pedrós & Ruiz-Trillo, 2014; King, 2004; King *et al.*, 2003; Miller, 2012).Because of how cells arise in fungi, communication along and between hyphae necessarily follows different principles. There are well-understood mechanisms for information processing along hyphae in vegetative mycelia, whereas there is no functional analogue of plasmodesmata or gap junctions that would mediate crosstalk between neighbouring(Bloemendal & Kuck, 2013) hyphae in fruiting bodies. Intercellular communication in fungi relies on the diffusion of chemical signals through the extracellular space, such as pheromones, volatile compounds, quorum sensing molecules(Albuquerque & Casadevall, 2012; Cottier & Mühlschlegel, 2012; Wongsuk, Pumeesat & Luplertlop, 2016) or even small proteins(Gyawali, Upadhyay, Way *et al.*, 2017; Wang, Tian, Gyawali *et al.*, 2013). It has evolved to signal through a loosely occupied space or among unicells, that primarily and suits the needs of vegetative mycelium or yeast cells. Nonetheless, such systems could be easily co-opted to communicate across tightly arranged hyphae in fruiting bodies, as suggested by a higher diversity(Busch & Braus, 2007; Frey, Reschka & Poggeler, 2015; Kuck, Beier & Teichert, 2016; Pöggeler *et al.*, 2006; Stajich, Wilke, Ahren *et al.*, 2010) of certain kinase gene families in fruiting body forming fungi, the expression of several kinases in fungal fruiting bodies and defects in fruiting body development in many kinase mutants(Pöggeler *et al.*, 2006). Remarkably, defects in either of the three MAP kinase pathways of fungi (OS, CWI and PG-MAPK pathways) impact fruiting body initiation(Kicka & Silar, 2004; Lichius *et al.*, 2012). Although the precise mechanisms of interhyphal communication within fruiting bodies have remained unknown so far, the lack of intercellular channels between neighboring hyphae suggest that fungi use different strategies to orchestrate the functioning of complex multicellular structures compared to plants and animals.

## 7. Is there a large genomic hurdle to complex multicellularity?

Although a complete understanding of MC-related genetic elements is lacking for any lineage, the significant the increases in phenotypic complexity associated with the evolution of complex multicellularity suggested the necessity of a comparably large set of genetic novelties(Cock *et al.*, 2010; Knoll, 2011). It would also accord well with it being a rare event in evolution. Genetic innovations underpinning the evolution of multicellularity have mostly been discussed in the context of gene duplications(Brawley *et al.*, 2017; Brunet *et al.*, 2017; Cock *et al.*, 2010; Miller, 2012; Richter *et al.*, 2013; Rokas, 2008; Sebe-Pedros *et al.*, 2017; Stajich *et al.*, 2010) and to a lesser extent in other sources of genetic novelty. In fungi, a number of transitions to complex multicellularity are coupled with surprisingly limited gene family diversification. The genus *Neolecta* (Taphrinomycotina) and fruiting body forming members of the Tremellomycetes and Pucciniomycotina, possess small genomes with a secondarily reduced protein coding capacity, similar to that of secondarily unicellular yeasts(Nagy; Stajich *et al.*, 2010). Consistent with an independent origin of complex multicellularity, *Neolecta* is nested in a clade of yeast-like and simple multicellular fungi (Taphrinomycotina) (Fig 5), which, we estimate, split from its closest extant complex MC relative >500 million years ago (based on ref(Kohler *et al.*, 2015)). Yet, its genome encodes as few as 5500 protein-coding genes (fewer than that of *Saccharomyces)* and very limited gene family diversification has been inferred along the evolutionary route to *Neolecta(Nguyen et al., 2017).* This is consistent with three hypotheses. First, the genetic hurdle to complex multicellularity may not be big and it may be relatively 'easy' for fungi to evolve complex multicellular structures. Second, gene duplications might not be the key changes underlying the evolution of complex multicellularity. Rather, building on a conserved gene repertoire shared with other Ascomycota, other sources of genetic innovations (e.g. gene regulatory network rewiring, alternative splicing patterns, noncoding RNA species, etc..) could underlie the independent origin of fruiting bodies in *Neolecta,* similarly to the picture that started to emerge from studies of animal multicellularity(Grau-Bove, Torruella, Donachie *et al.*, 2017; Richter *et al.*, 2013; Sebe-Pedros *et al.*, 2017). Third, a single origin of fruiting bodies in the Ascomycota could explain the limited gene family diversification on the evolutionary path leading to *Neolecta,* but would not explain the lack of known fruiting body genes of the Pezizomycotina from its genome. Phylogenetically it would also be a quite unparsimonious scenario, requiring several losses of fruiting body production in the Taphrinomycotina and the Saccharomycotina, among others. Which of these, or their combination explain best the evolution of complex MC in *Neolecta* and other fungi, in general, remains to be understood.

## 8. How many origins of complex multicellularity in fungi?

Complex multicellularity in fungi is a typical patchy(Telford *et al.)* character that appears in many phylogenetically distant clades. The prevailing view is that fungal fruiting bodies arose through convergent evolution(Knoll, 2011; Schoch *et al.*, 2009; Sebe-Pedros *et al.*, 2017; Stajich *et al.*, 2009; Taylor *et al.*, 2010), which is supported by the apparent lack of homologies between fruiting bodies in different clades. We above discussed 8 major clades of complex multicellular fungi (Fig 1), although there might be as many as 12, depending on the number of independent fruiting body forming clades in the Pucciniomycotina. If all of these clades evolved complex multicellularity independently, it means that there are 8-12 origins of this trait within fungi, compared to only four outside fungi. The large number and density of complex multicellular clades, however, prompts us to examine alternative views on the origin of complex MC in fungi. How would models implying a single origin of complex MC compare to ones implying multiple origins? Phylogenetically, the multiple origins model is more parsimonious than the single origin model, requiring 8-12 origins compared to 1 origin and >16 losses to explain the distribution of complex multicellularity across fungi (Fig. 3). However, purely phylogenetic considerations have little power to evaluate evolutionary hypotheses as the likelihood of the recurrent evolution of multigenic traits might be orders of magnitudes lower than that of a single origin followed by multiple losses. Therefore, in the following section we discuss how the phylogenetic conservation of developmental modules, genes and pathways underlying fruiting body development fits alternative scenarios of the evolution of complex MC in fungi.

**Figure 3.**
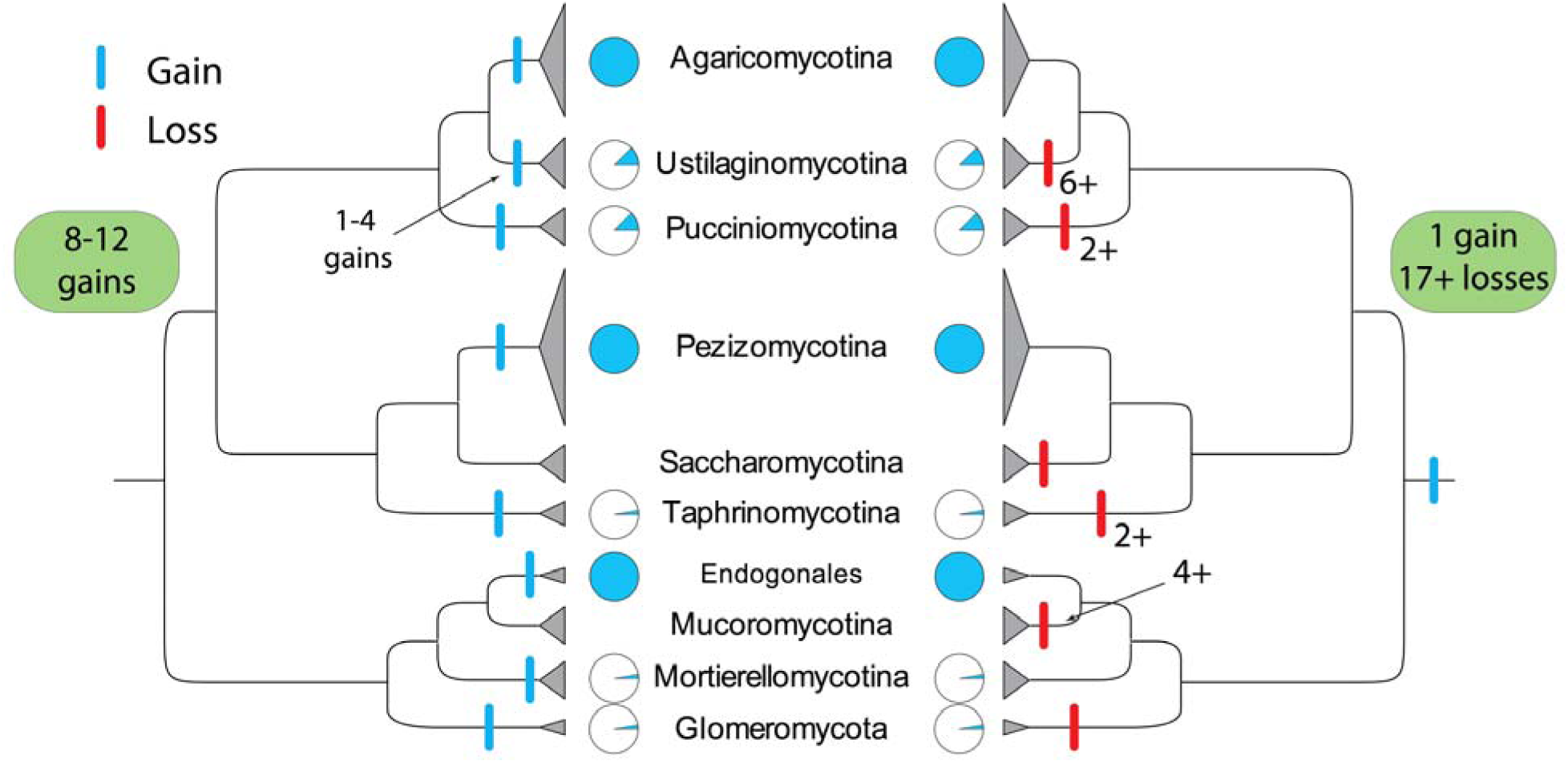
Alternative phylogenetic models for the recurrent origins of complex multicellularity in fungi. Gains and losses of complex multicellularity across fungi under two contrasting models are shown by vertical blue and red bars, respectively. Phylogenetically, the model implying convergence requires 8-12 independent origins to explain the phylogenetic distribution of complex multicellular fungi, whereas a model implying a single origin required 1 gain and >17 losses. Clades containing complex MC species are marked by pie charts with the blue section corresponding to the estimated fraction of complex multicellular species.

### 8.1 Homologies between independently evolved complex multicellular fungi?

If complex multicellular structures in disparate clades share homology, it should be detectable among genes involved in fruiting body development in the Asco- and Basidiomycota. Fruiting bodies in these clades show no evident homology at the phenotype level, however, this comes at no surprise as phenotypes can diverge quickly and so a more appropriate question is whether homologies exist at the level of the underlying genetic background. Some Asco- and Basidiomycota fruiting bodies comprise the best researched complex multicellular structures of fungi and the model species are as distant phylogenetically as any two complex multicellular clades are. In particular, are there homologies at the level of interactions high in the gene regulatory networks (including the initiation of the complex multicellular phase) and the key cellular functions of complex MC (adhesion, communication, development)?

The development of complex multicellular structures is part of the sexual reproductive program in fungi. In the most general sense, sexual reproduction, including mate detection, cell fusion and the formation of sexual propagules, and many of the associated genetic pathways are conserved across fungi. Fruiting bodies evolved to protect the developing sexual progeny and thus any gene regulatory network orchestrating their development should be plugged into the pathways governing sexual development. Indeed, mating genes regulate several aspects of fruiting body development: the formation of fruiting body initials (protoperithecia) of the Ascomycota, and that of primary and secondary hyphal knots of *Coprinopsis cinerea* are regulated by the *A* and *B* mating-type genes(Kues, Granado, Hermann *et al.*, 1998; Kues, Walser, Klaus *et al.*, 2002). On the other hand, protoperithecia of *Neurospora crassa* appear in a mating-independent manner, before fertilisation by a conidium of opposite mating type happens. Both protoperithecium and primary hyphal knot formation are induced by nutrient (mostly N_2_) starvation(Pöggeler *et al.*, 2006) through mechanisms that are widely conserved across fungi(D’Souza & Heitman, 2001; López-Berges, Rispail, Prados-Rosales *et al.*, 2010; Shertz, Bastidas, Li *et al.*, 2010) (Fig. 4) and even deeper in the eukaryotes (*e.g. Dictyostelium(Dubravcic, van Baalen & Nizak, 2014)*). More generally, nutrient availability is an important signal for sex in fungi: nutrient sensing pathways regulate sexual development through the mating type genes(Lengeler, Davidson, D’Souza *et al.*, 2000), similarly to many other processes that impact fruiting body development.

**Figure 4.**
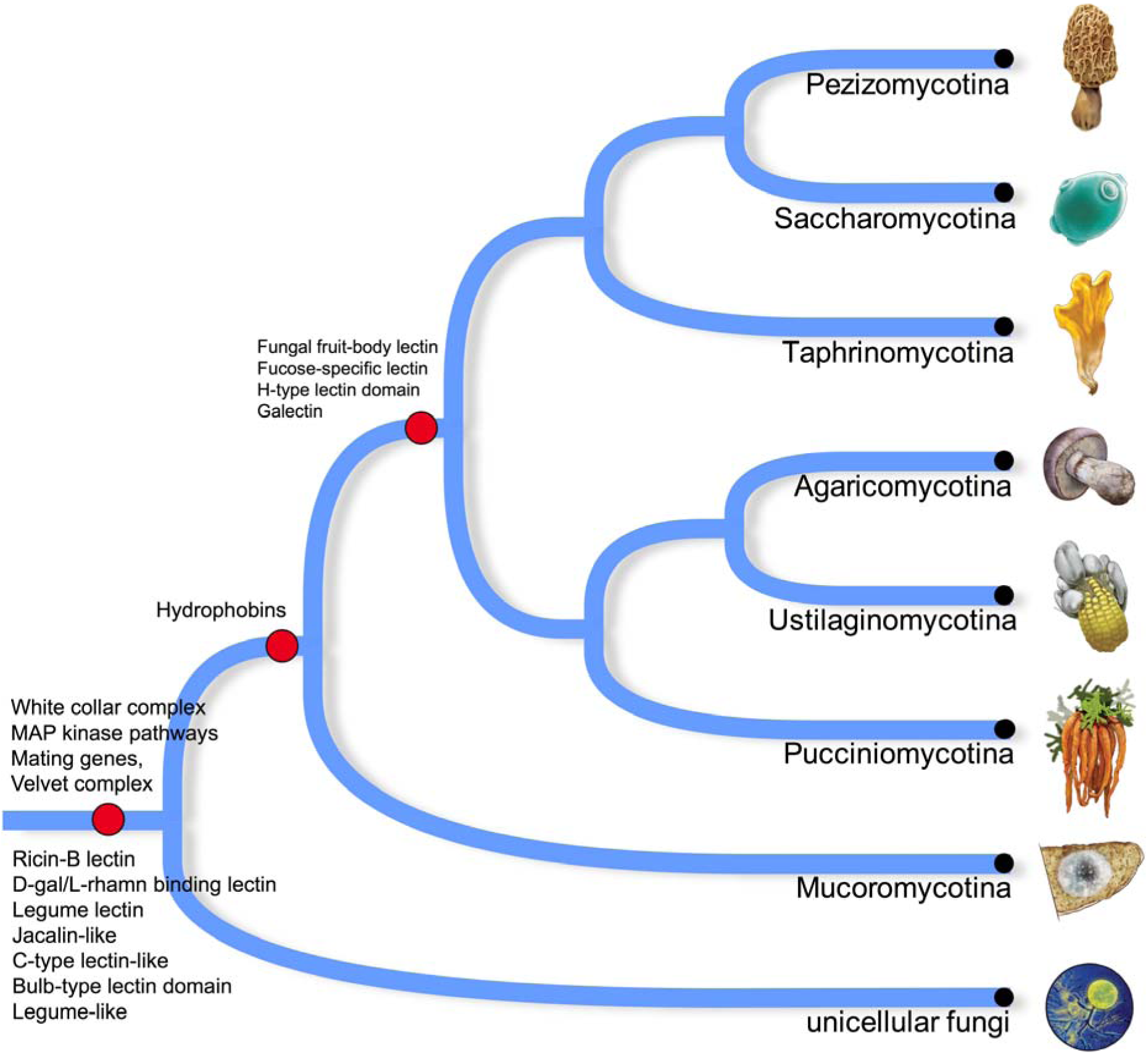
The conservation of characteristic gene families related to complex multicellularity in fungi. Several gene families involved in cell adhesion, defense, fruiting body initiation and morphogenesis are conserved across fungi, suggesting that the genetic prerequisites for multicellular functioning are widely available in uni- and simple multicellular fungi. Note that the emergence of most families predates the divergence of major clades of complex multicellular fungi, including the largest clades Pezizomycotina and Agaricomycotina.

The initiation of the complex multicellular phase is dependent on a number of additional factors, such as changing environmental variables (e.g. temperature, CO_2_ concentration) and the perception of external signals, such as light, by the vegetative mycelium. Light sensing relays several important processes of fruiting body development, including its initiation, maneuvering growth into the right direction and sensing seasonal light/dark periodicity that triggers fruiting(Kamada *et al.*, 2010; Pöggeler *et al.*, 2006). Many of these responses are mediated by the blue light receptor white collar complex (WCC), which, including its regulatory role in fruiting body development is conserved widely(Idnurm & Heitman, 2005; Rodriguez-Romero, Hedtke, Kastner *et al.*, 2010; Verma & Idnurm, 2013), although the specific interaction may differ even between closely related species(Kim, Kim, Lee *et al.*, 2015; Purschwitz, Müller, Kastner *et al.*, 2008). The WCC complex regulates sexual reproduction through mating genes(Idnurm *et al.*, 2005) in all fungal species examined so far(Idnurm & Heitman, 2010), except for budding and fission yeasts in which the complex has been lost(Nguyen *et al.*, 2017).

Similarly, the gross structure of mating pathways, that of mating loci and the regulation of sexual reproduction by mating genes is conserved across the Dikarya (Asco- and Basidiomycota) and maybe even earlier fungi(Casselton, 2002; Jones & Bennett, 2011; Kim, Wright, Park *et al.*, 2012; Raudaskoski & Kothe, 2010). G-proteins and the mitogen-activated protein kinase cascade that transduces the signal of compatible mate partner to the nucleus are also highly conserved across fungi(Ait Benkhali, Coppin, Brun *et al.*, 2013; Jones *et al.*, 2011; Kruzel, Giles & Hull, 2012), although differences between species exist at the level of terminal transcription factor identity(Kruzel *et al.*, 2012). Other two MAP kinase pathways (cell wall integrity and osmoregulatory) are also highly conserved across fungi and required for fruiting body development(Lichius *et al.*, 2012). The velvet complex coordinates differentiation processes and influences (a)sexual fruiting body development. Velvet complex proteins originated before the last common ancestor of complex multicellular lineages and are conserved across most fungi(Bayram & Braus, 2012) (Fig. 4).

On the other hand, little is known about the conservation of effector genes and cellular differentiation pathways (e.g. genes involved in morphogenesis, differentiation, etc.) that implement the complex multicellular phase. Adhesion-related GPI anchored proteins as well as hydrophobins are conserved across all fungi and are involved in fruiting body development in both the Asco- and Basidiomycota(Bruneau, Magnin, Tagat *et al.*, 2001; Costachel, Coddeville, Latge *et al.*, 2005; Robledo-Briones & Ruiz-Herrera, 2013; Szeto, Leung & Kwan, 2007). However, given their different roles in simple multicellular and yeast species, whether their widespread role in fruiting body development evolved through parallel co-option or reflects a plesiomorphic condition is difficult to decide. Similarly, lectins have been implicated in adhesion and defense in both the Asco- and Basidiomycota fruiting bodies(Hassan *et al.*, 2015), although different clades (and often different species) made use of different lectin families.

The lack of discernible homology among known genetic aspects downstream of fruiting body initiation implies extensive convergence. This is underpinned by the fact that most transcription factors known to be involved in fruiting body morphogenesis are specific to either the Asco- or Basidiomycota, although conservation of function in sexual reproduction has been reported at the family level (e.g. HMG-box TF_-s_)(Ait Benkhali *et al.*, 2013), which might suggest a plesiomorphic role on cell differentiation or that certain functions tend to be recruited repeatedly for fruiting body development.

Taken together, the genetic toolkit of fruiting body development includes both universally conserved and lineage-specific elements, which suggests it has been assembled gradually during evolution. Whereas many aspects of fruiting body development show convergence, homology exists among regulatory gene circuits underlying the initiation of fruiting body development and might exist at the level of certain multicellular functionalities. This points to a single origin of some of the foundations of complex multicellularity in fungi, which is remarkable from the perspective of independent origins and raises the question of how conservation can be reconciled with genetic theories of convergent evolution.

Explaining phenotypic convergence is a major challenge in evolutionary biology. Convergence in the classic sense implies the lack of homology, although recent advances revealed that this concept does not hold for several convergently evolved traits and suggests that a more detailed view on evolutionary convergence is necessary(Gompel & Prud'homme, 2009; Nagy *et al.*, 2014; Prud'homme, Gompel & Carroll, 2007; Stern, 2013). Phenotypic convergence can arise as a result of a range of genetic processes that include contributions of both homology and homoplasy(Nagy *et al.*, 2014; Panganiban, Irvine, Lowe *et al.*, 1997; Shubin, Tabin & Carroll, 2009; Wake, Wake & Specht, 2011) (convergence/parallelism). Fruiting body formation is a complex developmental process, and its current manifestations in both the Asco-and Basidiomycota probably evolved in a gradual manner. Similarly, gene regulatory circuits that orchestrate fruiting body development certainly also evolved in a stepwise manner, building on ancient regulatory modules, but also on co-option of conserved genes and the evolution of new ones (Fig. 5). At the moment relevant information to precisely reconstruct the evolution of the genetic toolkits underlying Asco-and Basidiomycota fruiting bodies and to answer the question whether their ancestor (or even earlier ones) was capable of forming simple fruiting bodies is lacking. Nevertheless, the high phylogenetic density of complex MC clades and the conservation of some mechanisms on fruiting body development suggests that convergence in the strict sense may not adequately explain the evolution of complex MC in fungi.

**Figure 5.**
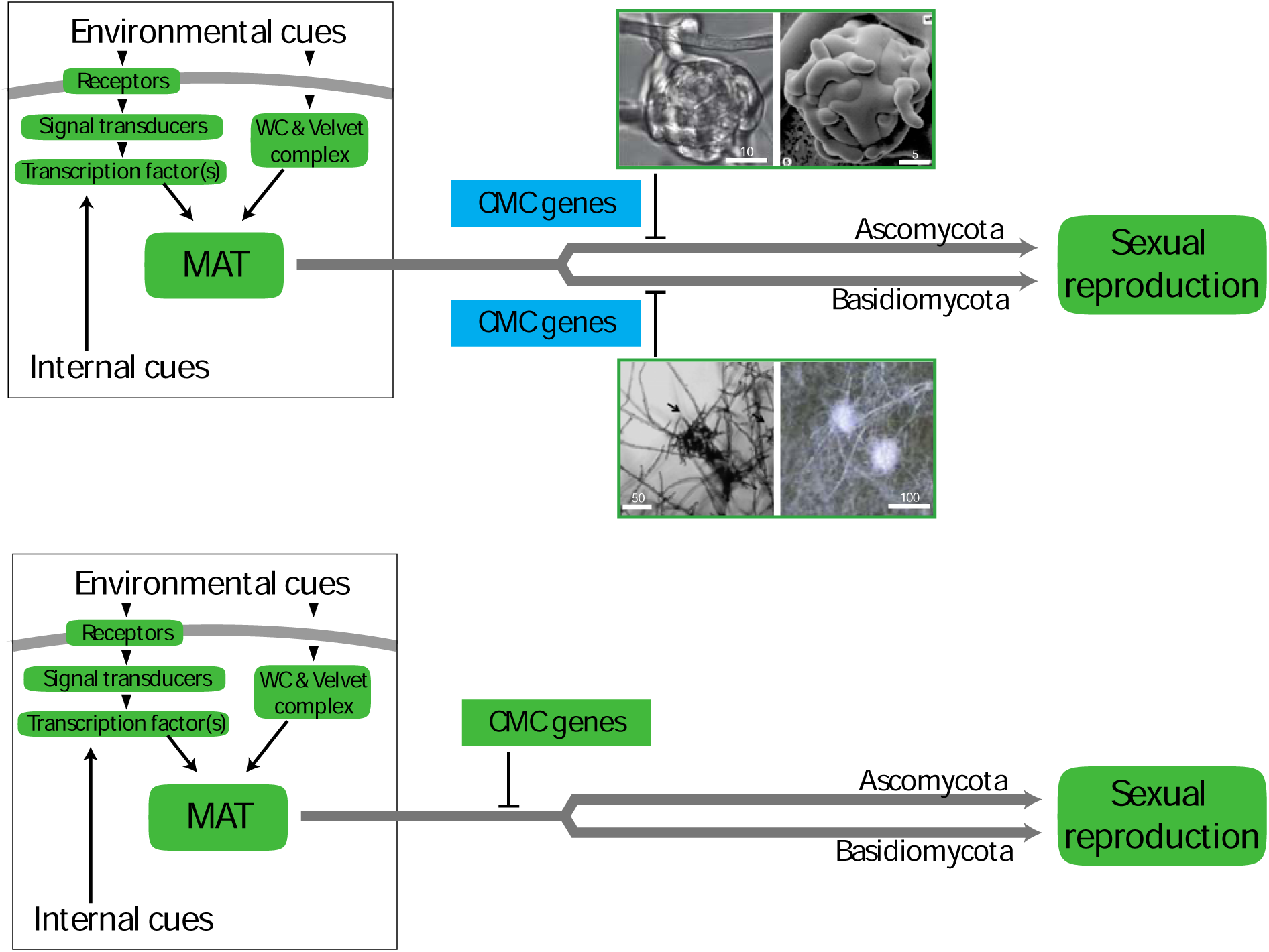
Two major alternative hypotheses for the evolution of complex multicellularity in fungi illustrated using a simplified case comprising Asco- and Basidiomycota. The initiation and trajectory of sexual reproduction in fungi comprises universally conserved mechanisms (highlighted in green). Genetic circuits involved in the development of fruiting bodies therefore should be linked into these conserved developmental pathways. A central question from the perspective of the evolution of fungal multicellularity is how genetic mechanisms of fruiting body development are linked to conserved circuits of sexual reproduction. The convergent origins model (top) implies that genetic mechanisms for fruiting body morphogenesis evolved independently along all lineages of complex multicellular fungi, whereas a single origin model (bottom) implies that at least part of the genetic toolkit of fruiting body development arose before the divergence of complex multicellular lineages. The presence of such genetic circuitries may predispose fungi for recurrently evolving complex multicellularity. The earliest complex multicellular stages, protoperithecia and primary hyphal knots for the Asco- and Basidiomycota are shown on the top image, respectively. Image sources are from Lord and Read (2010), Mayrhofer et al (2006), Lakkireddy et al (2011).

Understanding the components and conservation of early developmental modules that physically implement complex MC, downstream of the initiation of fruiting body development, thus, in our opinion, represents the key question for understanding the number of origins of complex multicellularity in fungi.

## Conclusions

(1) Fungi are one of the most enigmatic lineages of complex multicellular organisms. Although functional and mechanistic similarities with plant and animal multicellularity exist, there are fundamental differences in the driving forces, the timing and mechanisms of the evolution of simple and complex multicellularity in fungi, suggesting that there might be no unifying framework for the evolution of multicellularity across the tree of life. Is it possible then to establish general principles of the evolution of multicellularity? In terms of complex MC, there is certainly a common syndrome of traits that distinguish complex from simple multicellularity. This includes 3-dimensional organization, cell adhesion and an integrated developmental program that results in a multicellular structure or individual with genetically determined size and shape. For most lineages, complex MC comprises the reproducing individual, whereas it serves mostly reproductive roles in fungi. This is a fundamental difference between fungi and other lineages and provides an adaptive explanation for the patchy phylogenetic distribution of complex multicellularity in fungi.

(2) Complex multicellular fungi fall into 8-12 clades. This recurrence is currently considered to have happened through convergent evolution. While the genetic bases of several key aspects (e.g. morphological) of complex MC are lineage-specific and thus likely evolved convergently, most mechanisms of fruiting body initiation are universally conserved and thus likely have a single origin in fungi. How did morphogenetic processes that link the conserved and lineage-specific developmental modules evolve is among the least known aspects of fruiting body development currently, yet these might represent the crux of the matter for understanding the origins of complex multicellularity in fungi. Whether a single or multiple origins can explain the patchy phylogenetic distribution of complex multicellularity in fungi will need further research and we conjecture that focusing on the earliest cell-differentiation events in the development of complex multicellular structure holds the key to answering this question.

(3) Complex MC can be encoded by very small, yeast-like genomes, suggesting that complex MC does not require a great deal more genes than the development of simple multicellular fungi or yeats. Protein coding repertoires of fungal genomes fail to adequately explain differences in complexity level, and call for assays of other sources of genetic innovations(Nagy), including gene regulatory network rewiring, alternative splicing, various non-coding RNA species or RNA-editing pathways(Teichert, Dahlmann, Kuck *et al.*, 2017). Uncovering the genetic underpinnings of the evolution of complex MC in fungi is key to understanding the general principles of evolution towards increasingly more complex organisms. Our views on evolutionary trends towards complex MC in the tree of life and whether it represents a major transition in terms of genetic novelty hinges to a large extent on what we have learned and are about to learn through fungi. We expect that the unique ways of fungi for multicellular functioning could change paradigms in one of the central questions in biology.

## Acknowledgements

The authors appreciate fruitful discussions with Gábor M. Kovács on questions related to this review. The authors were supported by the Momentum Program of the Hungarian Academy of Sciences (LP2014/12) and ERC_HU Grant #118722 from the NRDI Office.

